# Are flowers tuned to buzzing pollinators? Variation in the natural frequency of stamens with different morphologies and its relationship to bee vibrations

**DOI:** 10.1101/2020.05.19.104422

**Authors:** Carlos Eduardo Pereira Nunes, Lucy Nevard, Fernando Montealegre-Zapata, Mario Vallejo-Marin

**Affiliations:** Department of Biological and Environmental Sciences. University of Stirling. FK9 4LA. Stirling, Scotland, United Kingdom; School of Life Sciences. University of Lincoln. LN6 7DL. Lincoln, England, United Kingdom

**Keywords:** Biomechanics, bumble bee, buzz pollination, flower diversity, resonance, *Solanum*

## Abstract

During buzz pollination, bees use vibrations to remove pollen from flowers. Vibrations at the natural frequency of pollen-carrying stamens are amplified through resonance, resulting in higher-amplitude vibrations. Because pollen release depends on vibration amplitude, bees could increase pollen removal by vibrating at the natural frequency of stamens. Yet, few studies have characterized the natural frequencies of stamens and compared them to frequencies of buzz-pollinating bees. Here we use laser Doppler vibrometry to characterise natural frequencies of stamens of six buzz-pollinated *Solanum* taxa of contrasting stamen morphology. We also compare the fundamental frequency of bumblebee buzzes produced on two *Solanum* species with different natural frequencies. We found that stamen morphology and plant identity explain variation in natural frequency of stamens. Our results show that medium-sized pollinators, such as bumblebees, produce buzzes of frequencies higher than the natural frequency of most (5/6) of the *Solanum* species we studied. However, the observed natural frequency of *Solanum* stamens is at the low end of the range of frequencies produced by other buzz-pollinating bees. Thus, our findings suggest that in some buzz pollination interactions, but not others, stamen resonance may play a role in mediating pollen release.

## 1. Introduction

More than half of all bee species evolved the ability to vibrate to extract pollen from flowers [1] giving rise to the syndrome of buzz pollination [2,3]. Most buzz-pollinated flowers present evolutionarily derived morphologies in which pollen locked inside stamens is released through small pores (poricidal stamens) [2]. While buzzing flowers, bees hold stamens using their mandibles and legs and activate their thoracic muscles [4] (Figure 1A). Pollen release from poricidal stamens is a function of the vibration characteristics, mainly their amplitude [5–10], which in turn depends on characteristics of the bee, the coupling between bee and flower [11,12], and the vibrational properties of the stamen (anther and filament) [3,13,14]. One descriptor of vibrational properties is the natural frequency. Natural frequencies are the modes in which a physical object can vibrate given its dimensions and material properties, such as rigidity or stiffness [15,16]. These natural frequencies determine how much a structure can bend (displacement amplitude) when vibrated. The natural frequency (first mode of vibration), is the lowest frequency in which an object can be moved in a way to increase vibration amplitude. In principle, if the vibrations applied by bees occurred at the natural frequency of stamens, vibration amplitude would increase, resulting in higher pollen removal [8,17].

**Figure 1.**
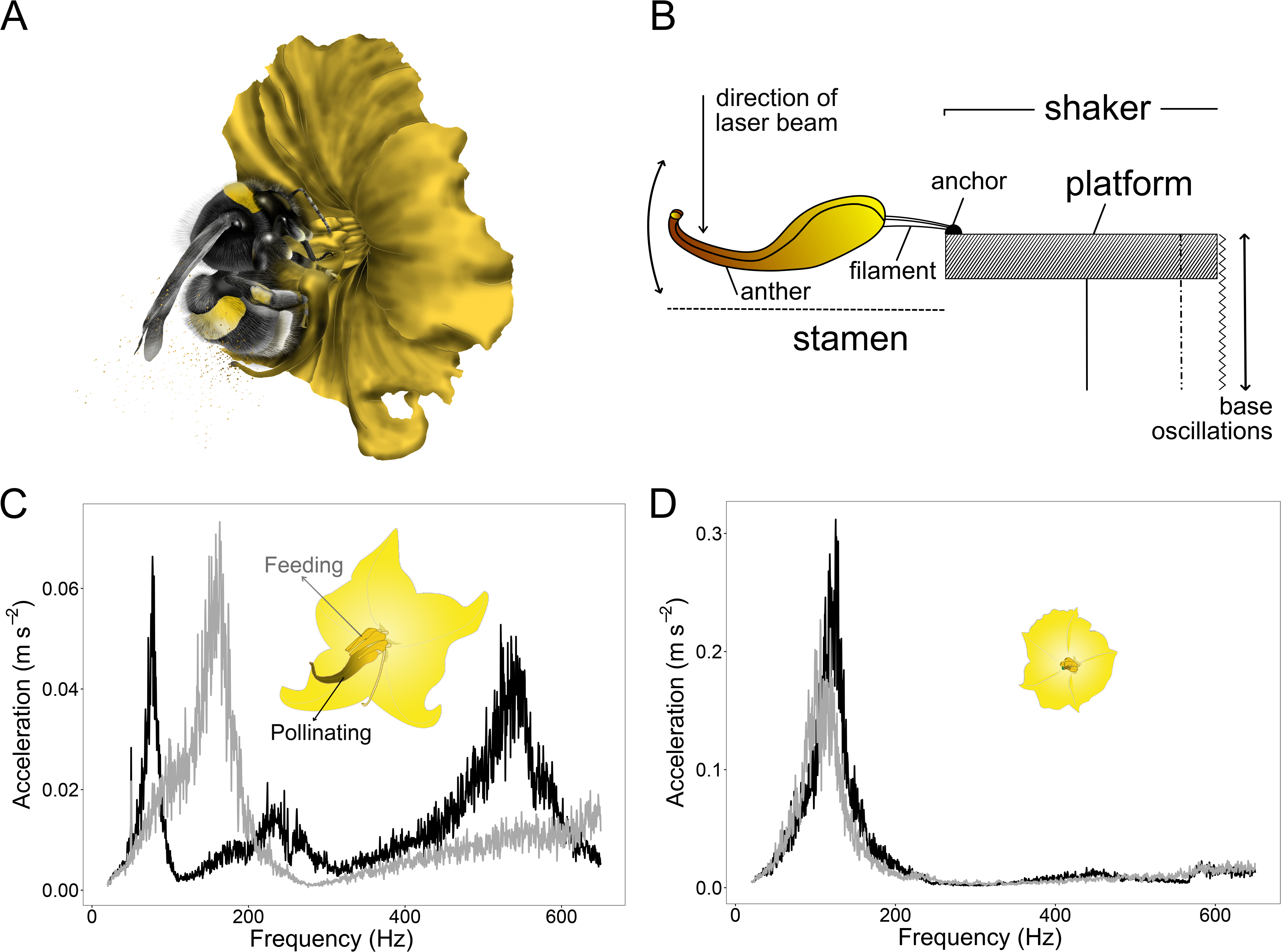
**(A.)** Illustration showing *B. terrestris* vibrating *S. rostratum*. **(B.)** Diagram of the setup in which stamens were attached to a platform on the magnetic shaker. **(C.)** Resonance pattern under white-noise vibration of feeding (grey line) and pollinating (black line) stamens of the heterantherous *Solanum rostratum* and **(D.)** of the non-heterantherous species *Solanum fructo-tecto*.

To date, little is known about the natural frequencies of stamens of buzz-pollinated plants. King and Buchmann [8] studied *Solanum laciniatum* and found that the stamen natural frequency (124 Hz) was significantly lower than the fundamental frequencies of bees buzzing these flowers (316 Hz). Other studies on natural frequencies of flowers have focused on wind-pollinated plants [17–19], in which vibrations induced by air flow lead to pollen ejection [8,20]. Here, we exploit natural variation both between and within plant species, to investigate the natural frequency of buzz-pollinated flowers. We use an unusual group of *Solanum* (Solanaceae) species that captures repeated independent transitions in flower and stamen morphology [21,22]. Unlike most *Solanum* species [23], species in section *Androceras* are heterantherous, bearing two sets of stamens with different morphologies specialised on either attracting and rewarding pollinators (feeding stamens) or fertilisation (pollinating stamens) [24,25]. We study three pairs of closely related taxa in which one member is large-flowered and highly heterantherous while the other is small-flowered and less heterantherous [21,26,27]. This combination of within-flower variation in stamen morphology and phylogenetically independent floral morphology transitions provides an unparalleled opportunity to investigate variation in natural frequencies in buzz-pollinated flowers. Our study addresses two questions: (1) To what extent do stamens with different morphologies have different natural frequencies? (2) Do bumblebees dynamically adjust the frequency of their vibrations while visiting flowers that differ in the natural frequency of their stamens? We predict that, all else being equal, longer stamens should have lower natural frequencies as predicted by theory of cantilever beams, and that bees should shift buzz frequency to match the visited flower, as this would increase pollen release.

## 2. Methods

### (a) Study systems

Six taxa of *Solanum* section *Androceras* (Solanaceae) native to Mexico and the southern USA were collected in the field (Table S1), and grown in the glasshouse as described in [21]. Vibration measurements were done in the lab (21°C). We used a single stamen cut at the base of the filament and measured two stamens from each flower (one feeding and one pollinating).

We used one colony of *Bombus terrestris audax* (Biobest, Westerlo, Belgium, Figure 1A) maintained according to [12]. We attached the colony to a flight arena (122×100×37 cm), illuminated with an LED light panel (59.5×59.5 cm, 48W Daylight; Opus Lighting Technology, Birmingham, UK) and kept on a 12:12h supplemental dark:light cycle. Room temperature was 22-23°C and humidity was 50-60% RH.

### (b) Data acquisition

#### Natural frequency of stamens

To measure natural frequencies we used a white noise playback approach [8,18]. Single stamens were exposed to white noise (20—20,000 Hz) generated in *Audacity* (Audacity Team 2019), using a linear power amplifier (LDS-LPA100, Brüel & Kjær, Nærum, Denmark) and a permanent magnetic shaker platform (LDV210, Brüel & Kjær). Each stamen was glued (Loctite Ultra Gel Control, Henkel, Hemel Hempstead, UK) by its filament base to a rigid platform at the top of the shaker (Figure 1B). We applied very low accelerations, at maximum 0.323 m s^−2^, which were not sufficient to remove pollen from flowers and ensured that the mass of the flower remained constant throughout each measurement.

To measure vibrations used a laser Doppler vibrometer (PDV-100, Polytec, Waldbronn, Germany) set to 500 mm s^−1^ sensitivity, a low-pass filter of 2500 Hz, and no high-pass filter. We focused the laser beam as close to the apical end of the stamen as possible at an axis perpendicular to the stamen length, parallel to the main axis of displacement of the shaker platform (Figure 1B). An accelerometer (0.8 grams, 352A24, PCB Piezotronics, Depew, NY) was attached to the shaker to record reference measurements. Both the laser vibrometer (recorded in acceleration units) and the accelerometer signals were simultaneously acquired using VibSoft-20 (Polytec) at a sampling frequency of 12,000 Hz, using a 20-5000 Hz bandpass filter, recorded for 1.28 s. From this, we obtained frequency spectra in the range from 20 to 2500 Hz using a fast Fourier transform (FFT; 6,400 lines) with a Hamming window (size=5,000) using VibSoft-20 and calculated the average frequency spectrum of 10 replicate samples for each stamen. Natural frequencies were estimated in an average of 10 flowers per taxon (range 5—11, n=53 flowers) from 2—8 individuals per taxon (two for *S. fructu-tecto*; average of 1.46 flowers per individual, n=41 individuals; Table S1). We measured eleven stamen and floral traits, including stamen length for the same 60 flowers to establish correlations among traits that could influence natural frequencies (Table S3). Preliminary analyses showed that all measured traits were positively correlated with each other, and negatively correlated with natural frequency (Figure S1). We therefore used only stamen length in subsequent analyses.

#### Frequency of bee vibrations

We used *S. citrullifolium* and *S. heterodoxum* to estimate the fundamental frequency of bee buzzes (Figure 2). We placed a single flower in the flight cage, allowing a bee to forage freely for approximately 10 minutes (visitation bout), and recording the audible component of floral vibrations (Zoom Hn4 Pro Handy, Zoom North America, Hauppauge, NY)[29] at 44kHz sampling rate for up to three minutes of floral buzzes. Fresh flowers were used for each bout. Naïve bees were first exposed to *S. citrullifolium* for six consecutive visitation bouts (n=10 bees), and buzzes in the first and sixth bout were recorded (n=1,640 buzzes analysed). Then, the same bees were exposed to *S. heterodoxum* for six additional bouts and buzzes in the first (n=10 bees) and sixth bouts (n=3 bees) recorded (n=758 buzzes). The lack of a reciprocal treatment (*S. heterodoxum*, then *S. citrullifolium*) reflects the reluctance of naïve *B. terrestris* to visit the small flowered taxon.

**Figure 2.**
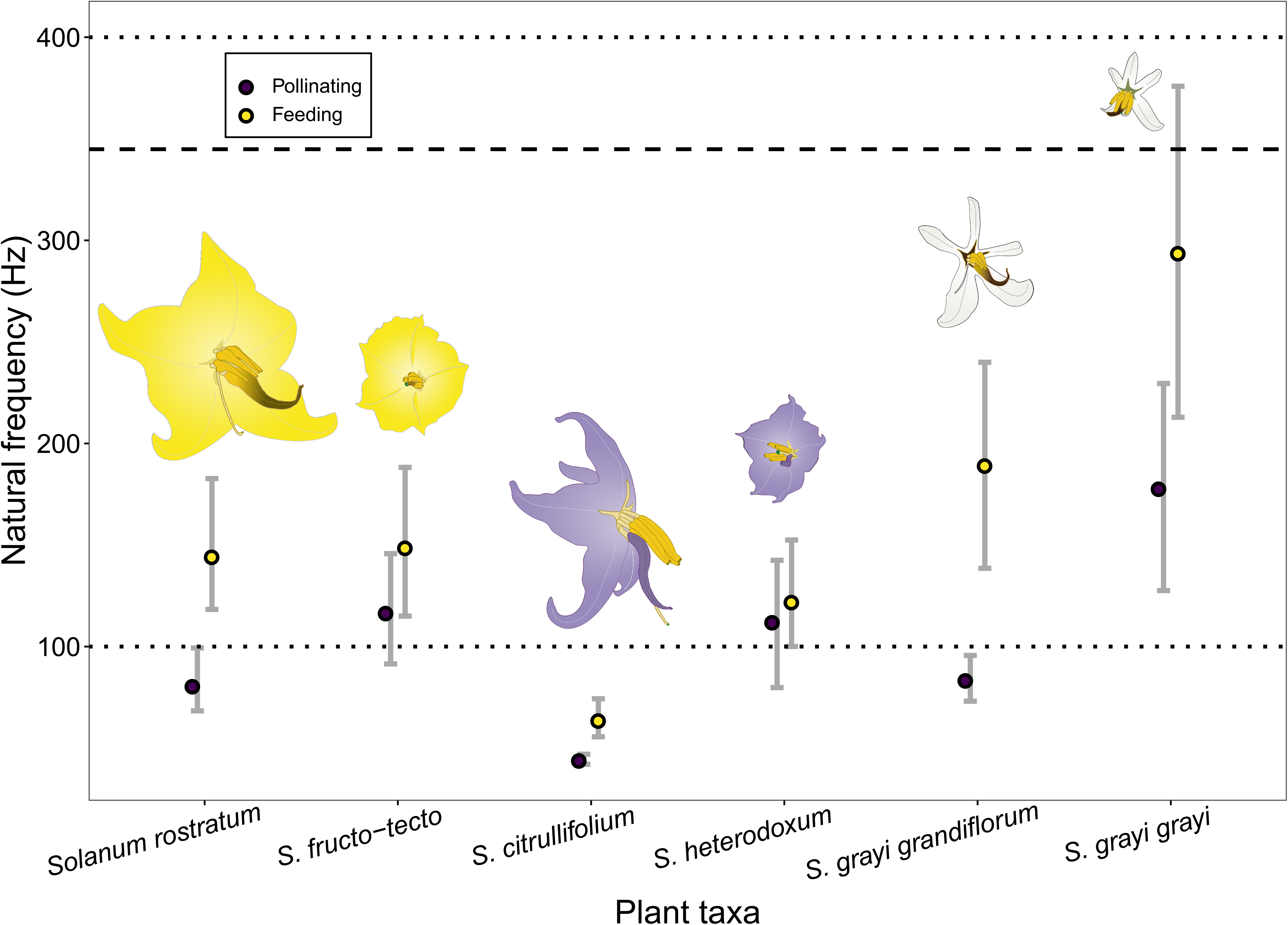
Natural frequencies of feeding and pollinating stamens of the six *Solanum* taxa studied here. The usual range of frequencies produced by buzz pollinating bees (100-400Hz) is shown with dotted lines, and the average frequency of *Bombus terrestris* on flowers of *S. citrullifolium* and *S. heterodoxum* obtained in the present study is shown with a dashed line. Flower illustrations depict variation in morphology and preserve size ratios among taxa.

### (c) Analyses

We visually identified the first (lowest) peak in the frequency spectrum (20-600 Hz range) and obtained its associated frequency [8,18]. To analyse variation in natural frequency and stamen characteristics, we fitted a linear mixed-effects model with natural frequency as the response variable, stamen length and stamen type as fixed effects, and plant taxon as a random effect using *lme4* [30]. Statistical significance of fixed effects was assessed with F-statistics with Satterwhite correction for degrees of freedom, implemented in *lmerTest* [31].

To obtain the fundamental frequency of bumblebees on *Solanum* flowers, we used *Audacity* to obtain the frequency spectrum (FFT) of each floral buzz using a Hamming window (size=512). To analyse the differences within bees’ fundamental frequencies, we fitted a linear mixed-effects model again using *lme4* with plant species and bout number as fixed effects, and individual bee identity as a random effect. We handled data with the help of *dplyr* [32], visualising it and creating plots with the help of *ggplot2* [33]. All analyses were done in *R* version 3.6.1 [34].

## 3. Results

### (a) Natural frequencies of stamens

The six *Solanum* taxa had natural frequencies between 43.75±6.15 Hz (mean ± SE) for pollinating stamens of *S. citrullifolium* and 294.30±47.37 for the feeding stamens of *S. grayi grayi* (Figure 2). Natural frequency differed among taxa and between stamen types, with pollinating stamens overall having lower natural frequencies than feeding stamens (Table 1, Table S2; Figure 2). Differences between stamen types were more marked in large flowered taxa (*S. rostratum, S. citrullifolium* and *S. grayi grandiflorum*), and smaller or absent in small flowered taxa (*S. fructu-tecto, S. heterodoxum*, and *S. grayi grayi*) (Figure 2). In each pair, pollinating stamens from the large-flowered taxon had lower natural frequencies than pollinating stamens from its small-flowered relative (Figure 2). After accounting for species identity and stamen type, we observed a marginally significant effect of stamen length on natural frequencies. Longer stamens tended to have lower natural frequencies than shorter stamens (Table1, Table S2).

**Table 1.** Effect of stamen type and length (mm) on the natural frequency of stamens from six taxa in *Solanum* section *Androceras* (Solanaceae). Model estimates and *p*-values were obtained using type III Sums of Squares of the fixed effects of a linear mixed-effects model. SE=Standard error.

| Model component | Estimate | SE | *p*-value |
| --- | --- | --- | --- |
| **Intercept** (feeding stamen) | 214.559 | 34.254 |  |
| **Stamen type** (pollinating stamen) | -37.013 | 13.712 | 0.008 |
| **Stamen length** (mm) | -7.598 | 3.755 | 0.055 |

### (b) Bee vibrations

We obtained 47-279 buzzes per bee per plant species (164±66 and 76±45 buzzes per bee, mean ± SD, for *S. citrullifolium* and *S. heterodoxum*, respectively). We found a small but statistically significant effect of both plant species and bout number on the fundamental frequency of floral buzzes (Table 2). Floral buzzes had a lower frequency in *S. heterodoxum* than in *S. citrullifolium*, and buzz frequency decreased with bout number (Table 2). *B. terrestris* vibrated flowers of *S. citrullifolium* at 345.25±0.87 Hz during their first bout (n=10 bees, 636 buzzes) and at 344.04±0.57 Hz in the sixth (n=10 bees, 1004 buzzes). The same individuals vibrated flowers of *S. heterodoxum* at 349.68±0.70 Hz (n=10 bees, 586 buzzes) and 329.47±0.95 Hz (n=3 bees, 163 buzzes), in their first and sixth bouts, respectively. In general, floral vibrations remained much higher than the natural frequency of the species being visited (Figure 2).

**Table 2.** Effect of plant species and bout number on the fundamental frequency (Hz) of floral vibrations produced by *Bombus terrestris* visiting flowers of two *Solanum spp*. Model estimates and *p*-values were obtained using type III Sums of Squares of the fixed effects of a linear mixed-effects model. SE=Standard error.

| Model component | Estimate | SE | *p*-value |
| --- | --- | --- | --- |
| **Intercept** (*S. citrullifolium*) | 348.734 | 3.397 |  |
| **Plant species** (*S. heterodoxum*) | -4.502 | 0.966 | 0.008 |
| **Bout number** | -0.733 | 0.150 | 0.002 |

## 4. Discussion

The natural frequencies of stamens of *Solanum* taxa studied here are within the range of frequencies achieved by bees during buzz pollination [4,9]. Consistently with previous work on bumblebees visiting mechanical flowers [35], our bumblebee experiment suggests that *B. terrestris* does not rapidly adjust their buzz frequency to match the natural frequency of the flowers they visit despite their ability to considerably change their buzz frequencies, e.g., between flower (313.09±2.63 Hz) and defence buzzes (236.32±4.29 Hz)[36]. The lack of matching could be explained by the fact that large pollinators like bumblebees can produce vibrations of sufficient force to expel pollen without needing the boost in vibration amplitude given by resonance, mostly relying on a “brute force” approach to remove pollen from anthers. Furthermore, stamen vibrational properties may not be the most important determinant of bee buzzes. Buzzing bees are hypothesised to use frequencies that correspond to the natural frequencies of their bodies to achieve higher accelerations while vibrating stamens, independent of the natural frequency of stamens or of the coupled bee-stamen system [7,12]. However, exploiting the vibrational properties of stamens may be more important for other bees unable to reach the required accelerations to elicit pollen release due to small size, mass, or biomechanical constraints [7]. Indeed, buzzing frequencies of small halictid bees range from 100 to 250 Hz and are more similar to the natural frequencies of *Solanum* stamens [8]. Further work should compare the stamen natural frequency of other buzz-pollinated flowers with the buzzing frequencies of the broader community of visiting bees to establish whether some bees can indeed exploit floral resonance for pollen release.

The variation in the natural frequency of stamens that we observed within and among closely related taxa suggests that this trait can evolve relatively rapidly. Within species, the lower natural frequencies of pollinating stamens compared to feeding stamens might be explained by morphology, geometry and material properties of stamen types [37] in addition to differences in length. Moreover, we find that the evolutionary transition to small flowers is reflected by changes in the vibrational properties of their stamens, particularly the degree of differentiation between pollinating and feeding stamens. Why feeding and pollinating stamens differ more strongly in large-than in small-flowered taxa remains unknown. It is possible that differences may allow flowers to regulate pollen release under a given buzz frequency. For example, a bee may be able to induce stamen resonance by matching its buzz natural frequency to that of the feeding stamens, but the same frequency will not induce resonance in pollinating stamens, potentially controlling pollen dispensing [10]. The convergence in natural frequency between stamen types observed in the small-flowered, less heterantherous taxa may reflect convergence of stamens within a flower into a single function. By building on classical work on the biomechanics of buzz-pollination [8] our work suggests new and exciting lines of inquiry integrating biomechanics and ecological interactions at the organismal level [38].

## Supporting information

Figure S1 - Correlation plot

Table S1 - Plant material accessions

Table S2 - Flower and bee frequencies

Table S3 - Morphological measurements

## Data accessibility

Data will be publicly deposited upon publication.

## Author’s contributions

CEPN carried out plant experiments, analysed the data, participated in study design and drafted the manuscript. LN carried out bee experiments, analysed the data, participated in study design and commented on the manuscript. FZM participated in experimental design and commented on the manuscript. MVM participated in conceiving the study and data analysis and helped draft the manuscript. All authors gave final approval for publication and agree to be held accountable for the work performed therein.

## Acknowledgements

We thank J. Kemp, D. Pritchard, D. Moore, and V. Brito for help with plant growth and discussions, and C. Woodrow for the bee illustration in Figure 1A.

## Supplementary Information

**Table S1.**□Information□on the origin of seeds□of the six taxa of□*Solanum*□Section□Androceras□studied here. Taxonomic classification follows Whalen (1979).□Names in bold for the more heterantherous taxa within each closely related pair.

**Table S2.** First resonant frequencies (Hz) for feeding and pollinating stamens from six *Solanum* taxa and peak frequencies emitted by bumblebees *Bombus terrestris audax* at flowers of *Solanum citrullifolium* and *S. heterodoxum*. Values in bold are significantly different when compared with pairwise Wilcoxon signed rank test for stamen types within each plant taxon and t-test for bee peak frequencies (p < 0.05).

**Table S3.** Morphological measurements of nine floral traits in the six *Solanum* taxa studied here.

**Figure S1.** Plot of the correlations among morphological variables accessed and the natural frequencies from stamens of the six *Solanum* taxa studied pooled.

