## Supplementary figures and images for "Are flowers tuned to buzzing pollinators? Variation in the natural frequency of stamens with different morphologies and its relationship to bee vibrations"

### Figure S1 - Correlation plot

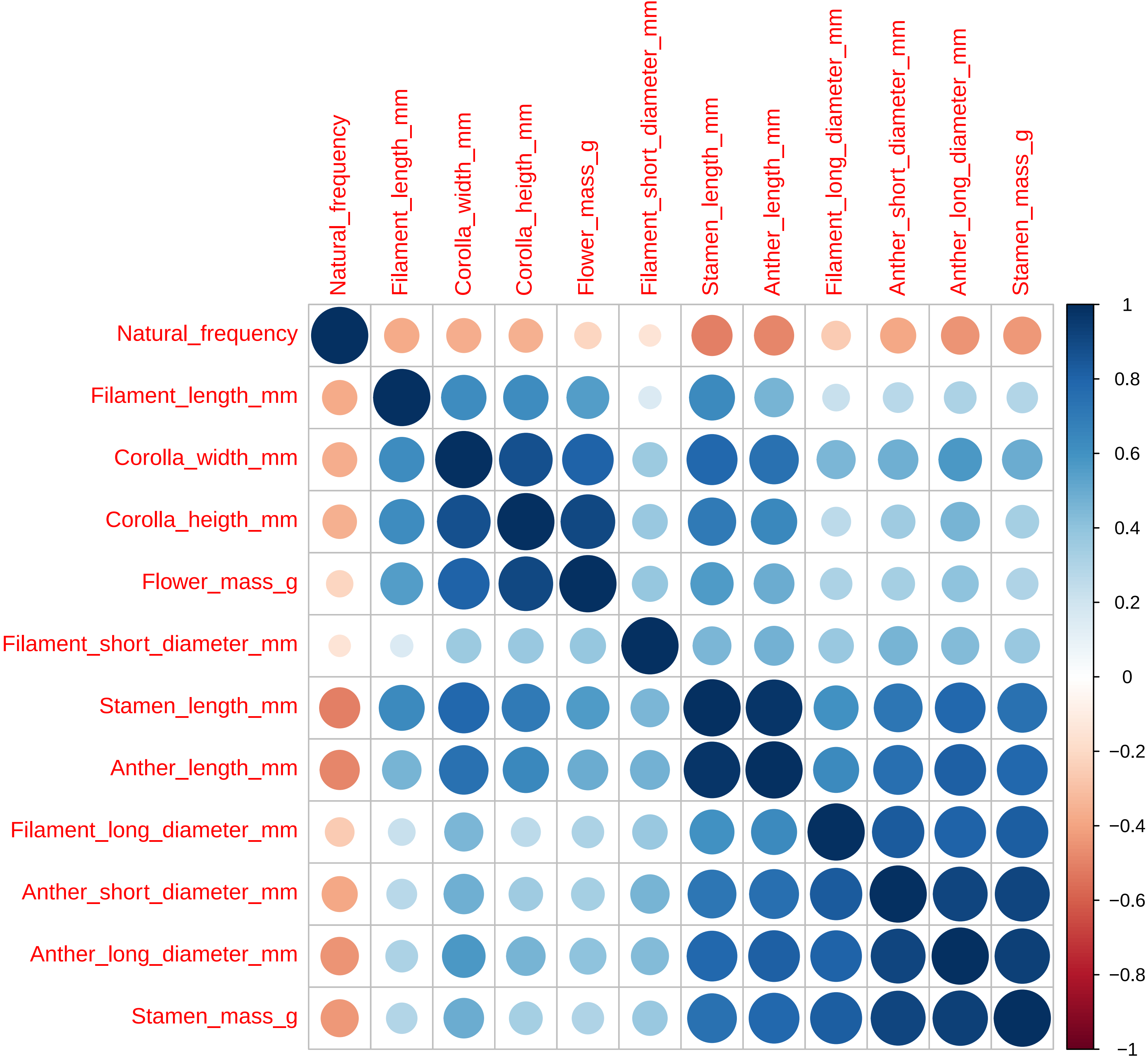
