## Supplementary material for "Are flowers tuned to buzzing pollinators? Variation in the natural frequency of stamens with different morphologies and its relationship to bee vibrations": Table S1 - Plant material accessions

**Table S1.** Information on the origin of seeds of the six taxa of *Solanum* Section Androceras studied here. Taxonomic classification follows Whalen (1979). Names in bold for the more heterantherous taxa within each closely related pair.

| **Taxon/Use** | **Accession** | **N of flowers** | **Total n of flowers** | **Localities** | **Latitude (N)** | **Longitude (W)** | **Elevation (m)** |
| --- | --- | --- | --- | --- | --- | --- | --- |
| Series Androceras |  |  |  |  |  |  |  |
| ***S. rostratum* Dunal** | 10-s-XX  10-s-79-23  10-s-79-24  10-s-82-8  10-s-77-13  10-s-77-5  10-s-77-12  10-s-79-25 | 1  2  1  1  1  2  2  1 | 11 | San Miguel, Querétaro | 20.90° | 100.71° | 2033 |
| *S. fructo-tecto* Cav. | 11-CU-4  10-AH-4 | 9  2 | 11 | Ciudad Universitaria, DF  Atitalaquia, Hidalgo | 19.39°  20.07° | 99.19°  99.22° | 2311  2090 |
| Series Pacificum |  |  |  |  |  |  |  |
| ***S. grayi* var. *grandiflorum* Whalen** | 08-s-78-9  08-s-78-6  08-s-78-24  08-s-79-10  08-s-79-19  08-s-79-21  08-s-79-20  08-s-78-6  08-s-79-18  08-s-78-12 | 1  1  1  1  1  1  2  1  1  1 | 11 | Tejupilco, Estado de México | 18.85° | 100.13° | 1375 |
| *S. grayi* var. *grayi* Whalen | 10-s196b-11  07-s-195b-23  07-s-194b-5  07-s-195b-12  07-s-194b-10  07-s-196b-24 | 2  1  1  1  1  1 | 7 | Los Álamos, Sonora | 27.00° | 108.93° | 385 |
| Series Violaceiflorum |  |  |  |  |  |  |  |
| ***S. citrullifolium* A. Braun** | 199-7-7-35  199-7-6-8  199-7-7-27  199-7-7-21  199-7-7-30  199-7-7-48  199-7-7-54 | 2  1  1  2  2  1  1 | 10 | Nijmegen, Solanaceae Collection | NA | NA | NA |
| *S. heterodoxum* Dunal | 11-PTEM-14-11  11-PTEM-15-13  11-PTEM-15-7  11-PTEM-14-8  11-PTEM-15-3  11-PTEM-14-9  11-PTEM-15-18  11-PTEM-14-4 | 1  1  1  1  1  3  1  1 | 10 | Teotihuacan Archaelogical Site, Estado de Mexico | 19.68° | 98.84° | 2284 |
