## Supplementary material for "Are flowers tuned to buzzing pollinators? Variation in the natural frequency of stamens with different morphologies and its relationship to bee vibrations": Table S2 - Flower and bee frequencies

**Table S2.** First resonant frequencies (Hz) for feeding and pollinating stamens from six *Solanum* taxa and peak frequencies emitted by bumblebees *Bombus terrestris audax* at flowers of *Solanum citrullifolium* and *S. heterodoxum*. Values in bold are significantly different when compared with pairwise Wilcoxon signed rank test for stamen types within each plant taxon and t test for bee peak frequencies (p < 0.05).

|  | **Stamen type** | | | |  |
| --- | --- | --- | --- | --- | --- |
| **Taxon** | **Feeding** | | **Pollinating** | |  |
|  | **Median ± i.q.r.** | **Mean ± s.e.** | **Median ± i.q.r.** | **Mean ± s.e.** | **N of flowers** |
| *Solanum rostratum* | **121.88 ± 41.21** | 144.85 ± 17.79 | **78.90 ± 17.38** | 81.14 ± 8.30 | 10 |
| *S. fructo-tecto* | 147.65 ± 78.91 | 149.40 ± 19.95 | 125.39 ± 43.36 | 117.97 ± 14.94 | 11 |
| *S. grayi* var. *grandiflorum* | **204.30 ± 112.21** | 189.77 ± 26.65 | **79.30 ± 22.27** | 80.89 ± 6.08 | 10 |
| *S. grayi* var. *grayi* | 310.55 ± 161.33 | 294.30 ± 47.37 | 200.00 ± 81.45 | 188.00 ± 30.76 | 5 |
| *S. citrullifolium* | **61.33 ± 14.65** | 64.22 ± 5.00 | **43.75 ± 6.15** | 44.57 ± 1.36 | 10 |
| *S. heterodoxum* | 109.77 ± 32.81 | 121.82 ± 14.65 | 124.61 ± 56.74 | 120.39 ± 16.92 | 7 |
|  | **Bee at flower mean ± s.e.** | | | | **N of bees (n of buzzes)** |
| *Bombus terrestris audax* at *S. citrullifolium* | 344.51 ± 0.49 | | | | 10 (1640) |
| *B. terrestris audax* at *S. heterodoxum* | 345.67 ± 0.68 | | | | 10 (758) |
