## Supplementary material for "Are flowers tuned to buzzing pollinators? Variation in the natural frequency of stamens with different morphologies and its relationship to bee vibrations": Table S3 - Morphological measurements

**Table S3.** Morphological measurements of nine floral traits in the six *Solanum* taxa studied here.

| **Taxon** | **Flower mass (g)** | **Pollinating stamen (PS) mass**  **(g)** | **Feeding stamen (FS) mass**  **(g)** | **PS length (mm)** | **FS length (mm)** | **PS filament major diameter (mm)** | **PS filament minor diameter**  **(mm)** | **FS filament major diameter (mm)** | **FS filament minor diameter (mm)** | **n** |
| --- | --- | --- | --- | --- | --- | --- | --- | --- | --- | --- |
| Series Androceras |  |  |  |  |  |  |  |  |  |  |
| *Solanum rostratum* Dunal | 0.118 ± 0.003 | 1.78E-02 ± 1.14E-03 | 3.45E-03 ± 1.81E-04 | 14.301 ± 0.177 | 9.584 ± 0.185 | 1.274 ± 0.047 | 0.542 ± 0.055 | 0.717 ± 0.031 | 0.464 ± 0.019 | 10 |
| *S*. *fructo-tecto* Cav. | 0.093 ± 0.006 | 1.42E-03 ± 6.86E-05 | 8.95E-04 ± 5.92E-05 | 6.150 ± 0.194 | 5.716 ± 0.232 | 0.685 ± 0.026 | 0.380 ± 0.020 | 0.487 ± 0.018 | 0.329 ± 0.013 | 11 |
| Series Pacificum |  |  |  |  |  |  |  |  |  |  |
| *S*. *grayi var. grandiflorum* Whalen | 0.06 ± 0.002 | 1.61E-02 ± 8.22E-04 | 2.03E-03 ± 7.72E-05 | 10.640 ± 0.199 | 7.486 ± 0.194 | 1.266 ± 0.045 | 0.427 ± 0.025 | 0.608 ± 0.031 | 0.350 ± 0.015 | 10 |
| S. *grayi var. grayi* Whalen | 0.028 ± 0.001 | 2.89E-03 ± 1.65E-04 | 1.43E-03 ± 1.03E-04 | 6.271 ± 0.145 | 5.403 ± 0.084 | 0.565 ± 0.029 | 0.332 ± 0.023 | 0.457 ± 0.028 | 0.315 ± 0.018 | 5 |
| Series Violaceiflorum |  |  |  |  |  |  |  |  |  |  |
| *S. citrullifolium* A. Braun | 0.086 ± 0.002 | 1.05E-02 ± 4.70E-04 | 4.24E-03 ± 2.30E-04 | 15.1065 ± 0.182 | 11.231 ± 0.233 | 0.841 ± 0.023 | 0.459 ± 0.012 | 0.603 ± 0.021 | 0.437 ± 0.017 | 10 |
| *S. heterodoxum* Dunal | 0.020 ± 0.001 | 4.66E-04 ± 3.52E-05 | 3.66E-04 ± 1.87E-05 | 4.259 ± 0.108 | 3.661 ± 0.102 | 0.408 ± 0.016 | 0.271 ± 0.017 | 0.319 ± 0.016 | 0.265 ± 0.013 | 7 |
